# Genome-wide association study implicates immune activation of multiple integrin genes in inflammatory bowel disease

**DOI:** 10.1101/058255

**Authors:** Katrina M. de Lange, Loukas Moutsianas, James C. Lee, Christopher A. Lamb, Yang Luo, Nicholas A. Kennedy, Luke Jostins, Daniel L. Rice, Javier Gutierrez-Achury, Sun-Gou Ji, Graham Heap, Elaine R. Nimmo, Cathryn Edwards, Paul Henderson, Craig Mowat, Jeremy Sanderson, Jack Satsangi, Alison Simmons, David C. Wilson, Mark Tremelling, Ailsa Hart, Christopher G. Mathew, William G. Newman, Miles Parkes, Charlie W. Lees, Holm Uhlig, Chris Hawkey, Natalie J. Prescott, Tariq Ahmad, John C. Mansfield, Carl A. Anderson, Jeffrey C. Barrett

**Affiliations:** Wellcome Trust Sanger Institute, Wellcome Trust Genome Campus, Hinxton, UK; Inflammatory Bowel Disease Research Group, Addenbrooke’s Hospital, Cambridge, UK; Institute of Cellular Medicine, Newcastle University, Newcastle upon Tyne; Division of Genetics and Rheumatology, Brigham and Women’s Hospital, Harvard Medical School, Boston, MA, USA; Program in Medical and Population Genetics, Broad Institute of Harvard and MIT, Cambridge, MA, USA; Precision Medicine Exeter, University of Exeter, Exeter, UK; IBD Pharmacogenetics, Royal Devon and Exeter Foundation Trust, Exeter, UK; Wellcome Trust Centre for Human Genetics, University of Oxford, Headington, UK; Christ Church, University of Oxford, St Aldates, UK; Gastrointestinal Unit, Wester General Hospital University of Edinburgh, Edinburgh, UK; Department of Gastroenterology, Torbay Hospital, Torbay, Devon, UK; Department of Child Life and Health, University of Edinburgh, Edinburgh, UK; Department of Paediatric Gastroenterology and Nutrition, Royal Hospital for Sick Children, Edinburgh, UK; Department of Medicine, Ninewells Hospital and Medical School, Dundee, UK; Guy’s & St Thomas’ NHS Foundation Trust, St Thomas’ Hospital, Department of Gastroenterology, London, UK; Translational Gastroenterology Unit, John Radcliffe Hospital, University of Oxford, Oxford OX3 9DS, UK; Human Immunology Unit, Weatherall Institute of Molecular Medicine, University of Oxford, Oxford OX3 9DS, UK; Paediatric Gastroenterology and Nutrition, Royal Hospital for Sick Children, Edinburgh, UK; Child Life and Health, University of Edinburgh, Edinburgh, Scotland, UK; Gastroenterology & General Medicine, Norfolk and Norwich University Hospital, Norwich, UK; Department of Medicine, St Mark’s Hospital, Harrow, Middlesex, UK; Department of Medical and Molecular Genetics, Faculty of Life Science and Medicine, King’s College London, Guy’s Hospital, London, UK; Sydney Brenner Institute for Molecular Bioscience, Faculty of Health Sciences, University of Witwatersrand, South Africa; Genetic Medicine, Manchester Academic Health Science Centre, Manchester, UK; The Manchester Centre for Genomic Medicine, University of Manchester, Manchester, UK; Translational Gastroenterology Unit and the Department of Paediatrics, University of Oxford, Oxford, United Kingdom; Nottingham Digestive Diseases Centre, Queens Medical Centre, Nottingham, UK; Institute of Human Genetics, Newcastle University, Newcastle upon Tyne, UK

## Abstract

Genetic association studies have identified 210 risk loci for inflammatory bowel disease^1–7^, which have revealed fundamental aspects of the molecular biology of the disease, including the roles of autophagy and Th17 cell signaling and development. We performed a genome-wide association study of 25,305 individuals, and meta-analyzed with published summary statistics, yielding a total sample size of 59,957 subjects. We identified 26 new genome-wide significant loci, three of which contain integrin genes that encode molecules in pathways identified as important therapeutic targets in inflammatory bowel disease. The associated variants are also correlated with expression changes in response to immune stimulus at two of these genes (*ITGA4, ITGB8*) and at two previously implicated integrin loci (*ITGAL, ICAM1*). In all four cases, the stimulus-dependent expression increasing allele also increases disease risk. We applied summary statistic fine-mapping and identified likely causal missense variants in the primary immune deficiency gene *PLCG2* and the negative regulator of inflammation, *SLAMF8*. Our results demonstrate that new common variant associations continue to identify genes and pathways of relevance to therapeutic target identification and prioritization.

---

Inflammatory bowel disease (IBD) is a chronic, debilitating, disorder of the gastrointestinal tract that includes two common disease subtypes, Crohn’s disease and ulcerative colitis. Disease pathogenesis is poorly understood but is likely driven by a dysregulated immune response to unknown environmental triggers in genetically susceptible individuals. Treatment regimes often use potent immunomodulators to achieve and maintain remission of symptoms. However, patients commonly experience side effects, lose response to treatment, or develop complications of IBD, with many ultimately requiring major abdominal surgery. Previous genome-wide association studies (GWAS) and targeted follow-up using the Immunochip have been very successful at identifying genetic risk loci for IBD, but increased biological understanding has not yet had a significant impact on therapy for these disorders.

In order to further expand our understanding of the biology of these disorders we carried out a GWAS of 12,160 IBD cases and 13,145 population controls of European ancestry that had not been included in any genome-wide meta-analysis of IBD to date (Supplementary Table 1, Online Methods). We imputed genotypes using a reference panel comprising whole genome sequences from 4,686 IBD cases^8^ and 6,285 publically available population controls^9,10^. Following quality control (Online Methods) we tested 9.7 million sites for association. At the 232 IBD associated SNPs in the latest meta-analysis by the International IBD Genetics Consortium^1^, 228 had effects in the same direction in our data, 188 showed at least nominal evidence of replication (P<0.05) and none showed significant evidence of heterogeneity of effect by Cochrane’s Q test. Among these replicated loci was a genome-wide significant association on chromosome 10q25 that was only previously significantly associated with Crohn’s disease in individuals of East Asian ancestry^3,7^, further supporting near complete sharing of genetic risk loci across populations^1^. We meta-analyzed our new GWAS data with previously published summary statistics from 12,882 IBD cases and 21,770 population controls imputed using the 1000 Genomes Project reference panel^1^ (Supplementary Figures 1-3, Supplementary Table 2). We observed inflation of the summary statistics (λ_GC_ = 1.23 and 1.29 for Crohn’s and ulcerative colitis, respectively), but LD score regression demonstrated that this was due to broad polygenic signal, rather than confounding population substructure (both intercepts = 1.09, Online Methods).

We identified 26 new loci at genome-wide significance (**Table 1**). In order to identify causal variants, genes and mechanisms, we performed a summary-statistic fine-mapping analysis on these loci, as well as 39 previously discovered loci where fine-mapping had not yet been attempted^11^ (Online Methods, Supplementary Table 3). In order to be confident about fine-mapping inferences, we restricted subsequent analyses to 12 signals where we had high quality imputed data for all relevant variants (Online Methods). At 6 of these 12 loci we identified a single variant with >50% probability of being causal (**Table 2**, Supplementary Figures 4-6). Among these were two loci where a single variant had >99% probability of being causal: a missense variant predicted to affect protein function in *SLAMF8*, (**Figure 1a**), and an intronic variant in the key regulator of Th17 cell differentiation, *RORC*^12^. SLAMF8 is a cell surface receptor that is expressed on activated myeloid cells and has been reported to negatively regulate inflammatory responses by repressing the production of reactive oxygen species (ROS)^13^ and inhibiting their migration to sites of inflammation^14^. *RORC* encodes RORγt, the master transcriptional regulator of Th17 cells^12^ and group 3 innate lymphoid cells^15^. Both of these cell types play important roles in defence at mucosal surfaces, especially in the intestine, and have been shown to contribute to the homeostasis between the intestinal immune system and gut microbiota^16,17^, an equilibrium that is known to be lost in inflammatory bowel disease^18^. Pharmacologic inhibition of RORγt has been shown to offer therapeutic benefit in mouse models of intestinal inflammation, and reduces the frequency of Th17 cells isolated from primary intestinal samples of IBD patients^19^.

**Figure 1.**
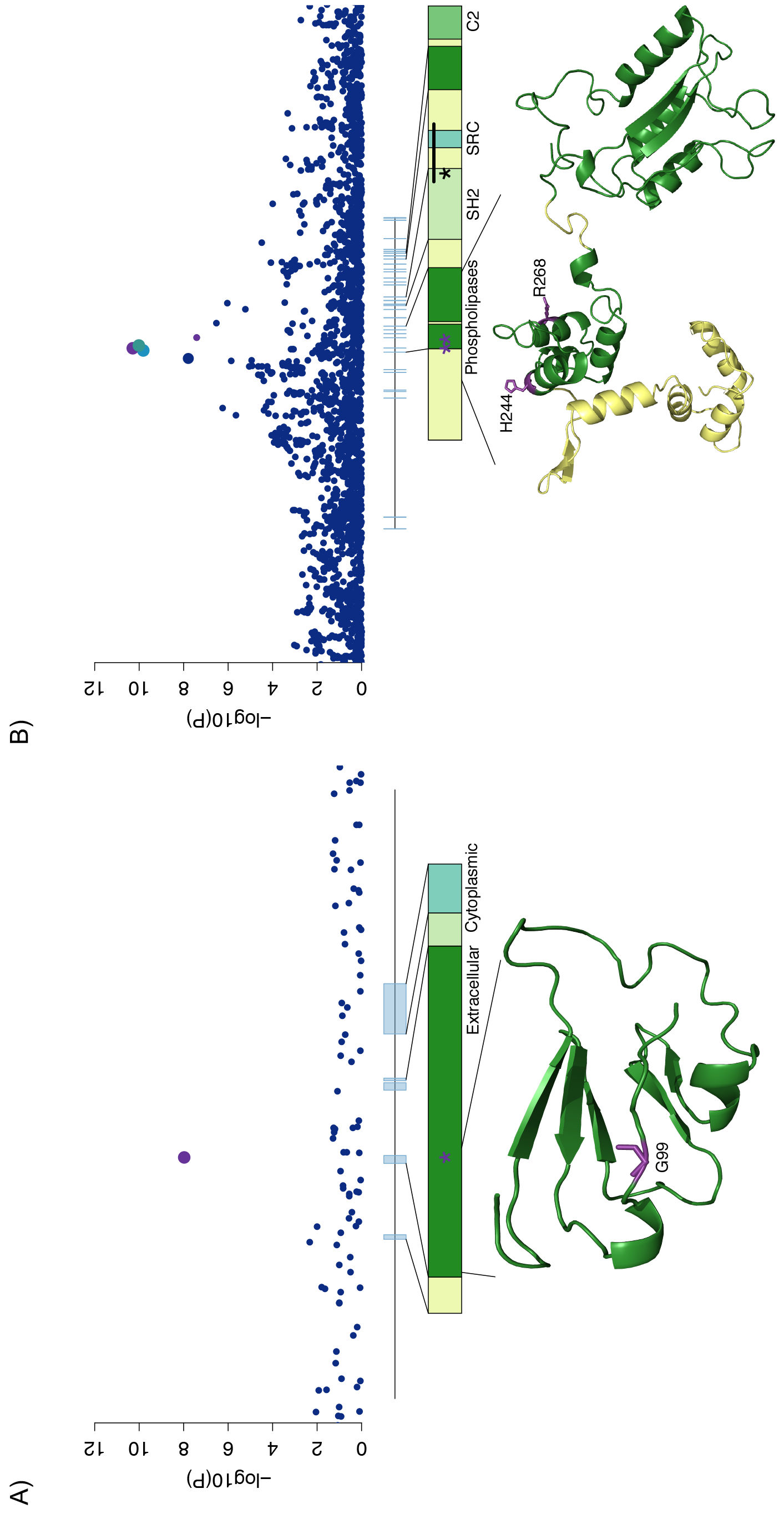
Likely causal missense variants. For A) SLAMF8 and B) PLCG2, local association results are plotted with point size corresponding to LD to our lead variant and color to fine-mapping probability (purple > 50%, intermediate blue 10-50%, navy blue <10%). Gene body diagrams and protein domain annotations are taken from ENSEMBL, and partial predicted crystal structures for both proteins are obtained from the SWISS-MODEL respository.

**Table 1.** Novel IBD-associated loci.

| Rsid | Chr | Position (bp) | Left-right (Mb) | Risk Allele | Non-risk Allele | Risk Allele Frequency in 1000 Genomes CEU+GBR | P<sub>Meta</sub> | OR (95% CI) | Phenotype |
| --- | --- | --- | --- | --- | --- | --- | --- | --- | --- |
| rs34687326 | 1 | 159799910 | 159.8-159.8 | G | A | 0.9 | 1.06 x 10<sup>-08</sup> | 1.18 (1.12-1.24) | CD |
| rs59043219 | 1 | 209970610 | 209.97-210.02 | A | G | 0.379 | 1.09 x 10<sup>-08</sup> | 1.08 (1.05-1.10) | IBD |
| rs6740847 | 2 | 182308352 | 182.31-182.33 | A | G | 0.508 | 1.22 x 10<sup>-13</sup> | 1.10 (1.07-1.12) | IBD |
| rs144344067 | 2 | 187576378 | 187.5-187.68 | A | AT | 0.895 | 1.29 x 10<sup>-08</sup> | 1.12 (1.08-1.16) | IBD |
| rs1811711 | 2 | 228670476 | 228.67-228.67 | C | G | 0.826 | 6.09 x 10<sup>-09</sup> | 1.14 (1.1-1.18) | UC |
| rs76527535 | 2 | 242484701 | 242.47-242.49 | C | T | 0.745 | 2.87 x 10<sup>-08</sup> | 1.09 (1.06-1.12) | IBD |
| rs2581828 | 3 | 53133149 | 53.1-53.17 | C | G | 0.597 | 6.46 x 10<sup>-09</sup> | 1.10 (1.07-1.13) | CD |
| rs2593855 | 3 | 71175495 | 71.16-71.19 | C | T | 0.663 | 2.54 x 10<sup>-09</sup> | 1.09 (1.06-1.11) | IBD |
| rs503734 | 3 | 101023748 | 100.91-101.27 | A | G | 0.513 | 2.67 x 10<sup>-08</sup> | 1.07 (1.05-1.10) | IBD |
| rs56116661 | 3 | 188401160 | 188.4-188.4 | C | T | 0.795 | 5.67 x 10<sup>-10</sup> | 1.14 (1.10-1.18) | CD |
| rs11734570 | 4 | 38588453 | 38.58-38.59 | A | G | 0.368 | 4.80 x 10<sup>-08</sup> | 1.07 (1.05-1.10) | IBD |
| rs17656349 | 5 | 149605994 | 149.59-149.63 | T | C | 0.466 | 1.54 x 10<sup>-08</sup> | 1.09 (1.06-1.13) | UC |
| rs113986290 | 6 | 19781009 | 19.72-19.83 | C | T | 0.989 | 7.59 x 10<sup>-09</sup> | 1.36 (1.25-1.46) | UC |
| rs67289879 | 6 | 42007403 | 42-42.01 | T | C | 0.179 | 3.04 x 10<sup>-08</sup> | 1.09 (1.06-1.13) | IBD |
| rs11768365 | 7 | 6545188 | 6.5-6.55 | A | G | 0.816 | 3.88 x 10<sup>-08</sup> | 1.09 (1.06-1.12) | IBD |
| rs149169037 | 7 | 20577298 | 20.58-20.58 | G | A | 0.895 | 3.26 x 10<sup>-08</sup> | 1.14 (1.10-1.19) | IBD |
| rs243505 | 7 | 148435339 | 148.4-148.58 | A | G | 0.624 | 3.04 x 10<sup>-10</sup> | 1.08 (1.06-1.11) | IBD |
| rs7911117 | 10 | 27179596 | 27.16-27.18 | T | G | 0.871 | 1.84 x 10<sup>-08</sup> | 1.14 (1.10-1.19) | UC |
| rs111456533 | 10 | 126439381 | 126.32-126.55 | G | A | 0.829 | 1.18 x 10<sup>-09</sup> | 1.11 (1.08-1.14) | IBD |
| rs11221335 | 11 | 128385906 | 128.38-128.4 | C | T | 0.224 | 2.44 x 10<sup>-08</sup> | 1.09 (1.06-1.12) | IBD |
| rs80244186 | 13 | 42917861 | 42.84-42.94 | C | T | 0.111 | 3.66 x 10<sup>-08</sup> | 1.13 (1.09-1.18) | CD |
| rs11548656 | 16 | 81916912 | 81.91-81.92 | A | G | 0.961 | 5.18 x 10<sup>-11</sup> | 1.27 (1.20-1.34) | IBD |
| rs10492862 | 16 | 82867456 | 82.87-82.92 | A | C | 0.308 | 1.26 x 10<sup>-09</sup> | 1.11 (1.08-1.15) | CD |
| rs4256018 | 20 | 6093889 | 6.08-6.1 | G | T | 0.25 | 1.23 x 10<sup>-08</sup> | 1.08 (1.05-1.11) | IBD |
| rs138788 | 22 | 35729721 | 35.72-35.74 | A | G | 0.418 | 2.95 x 10<sup>-08</sup> | 1.09 (1.06-1.13) | UC |
| rs4821544 | 22 | 37258503 | 37.26-37.26 | C | T | 0.321 | 1.76 x 10<sup>-08</sup> | 1.10 (1.07-1.13) | CD |

**Table 2.**
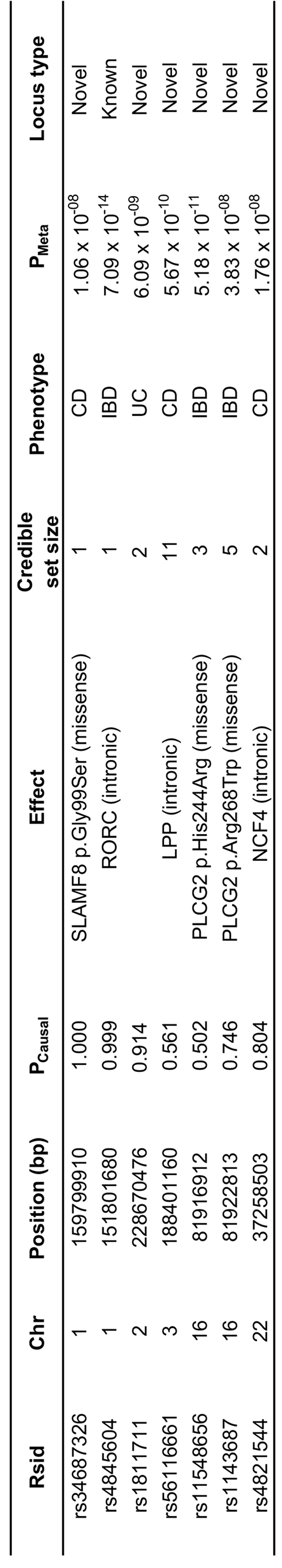
Variants fine-mapped to >50% probability of being causal in their given locus.

In loci where fine-mapping was less clearly resolved, we searched for likely functional variants, observing a missense variant predicted to affect protein function (CADD = 16.45, 50.2% probability of causality) in *PLCG2*. Furthermore, after conditioning on this variant, we discovered a second, independent, likely functional (CADD = 34, 74.6% probability of causality) missense variant in the same gene (*P*=2×10^−8^). *PLCG2* encodes a phospholipase enzyme that plays a critical role in regulating immune pathway signalling^20^, and has previously been implicated in two autosomal dominant immune disorders. Intragenic deletions in its autoinhibitory domain cause antibody deficiency and immune dysregulation (familial cold autoinflammatory syndrome 3, MIM 614468)^21^ and heterozygous missense variants (e.g. p.Ser707Tyr) lead to a phenotype that includes intestinal inflammation^22^ (**Figure 1b**).

A more general overlap between candidate IBD GWAS genes and Mendelian disorders of inflammation and immunity has been previously observed^23^. In addition to *PLCG2* we identified an association between Crohn’s disease and an intronic variant in *NCF4* (P=1.76 x 10^−8^). This gene encodes p40phox, a component of the NADPH-oxidase system that is responsible for the oxidative burst in innate immune cells and which is a key mechanism of killing phagocytosed bacteria. Rare pathogenic variants in *NCF4* cause autosomal recessive chronic granulomatous disease, characterized by Crohn’s disease-like intestinal inflammation and defective ROS production in neutrophils^24^. Our associated variant, rs4821544, had previously been suggestively associated with small bowel Crohn’s disease^25,26^, and when we stratified patients by disease location we found that the effect was consistently stronger for small bowel compared to large bowel disease (Supplementary Figure 7).

Among the remaining 22 novel loci we noted three that were within 150kb of integrin genes (***ITGA4***, ***ITGAV*** and ***ITGB8***), while two previously associated loci overlap with a fourth integrin, ***ITGAL***, and its binding partner ***ICAM1***. Integrins are cell adhesion mediators with bi-directional signalling capabilities that play a crucial role in leukocyte homing and cell differentiation in inflammation and cancer^27^. Given the strong candidacy of these genes, we sought potentially causal molecular mechanisms that would connect the IBD associated SNPs to integrin regulation. Our fine-mapping analysis excluded the possibility that these associations are caused by protein-coding changes, so we next tested for effects of IBD risk SNPs on integrin gene expression in immune cells using nine publicly available eQTL datasets. While many eQTLs and GWAS signals show some degree of correlation, inferences about causality require more robust statistical co-localization of the two signals. Remarkably, we observed three of our five associations had >90% probability of being driven by the same variants as monocyte-specific stimulus response eQTLs (***ITGA4***, ***P***_LPS_24hr_=0.984; ***ITGAL***, ***P***_LPS_24hr_=0.980; ***ICAM1***, ***P***_LPS_2hr_=0.961; Supplementary Table 4). A fourth association, ***ITGB8***, is difficult to map due to extended linkage disequilibrium in the locus, but shows intermediate evidence of co-localization (**P**_LPS_24hr_=0.712) in response to the same stimulus (**Figure 2**). These observations suggest upregulation of pro-inflammatory cell surface markers as a potential mechanism of action, as all four of the IBD risk increasing alleles upregulate expression of their respective genes.

**Figure 2.**
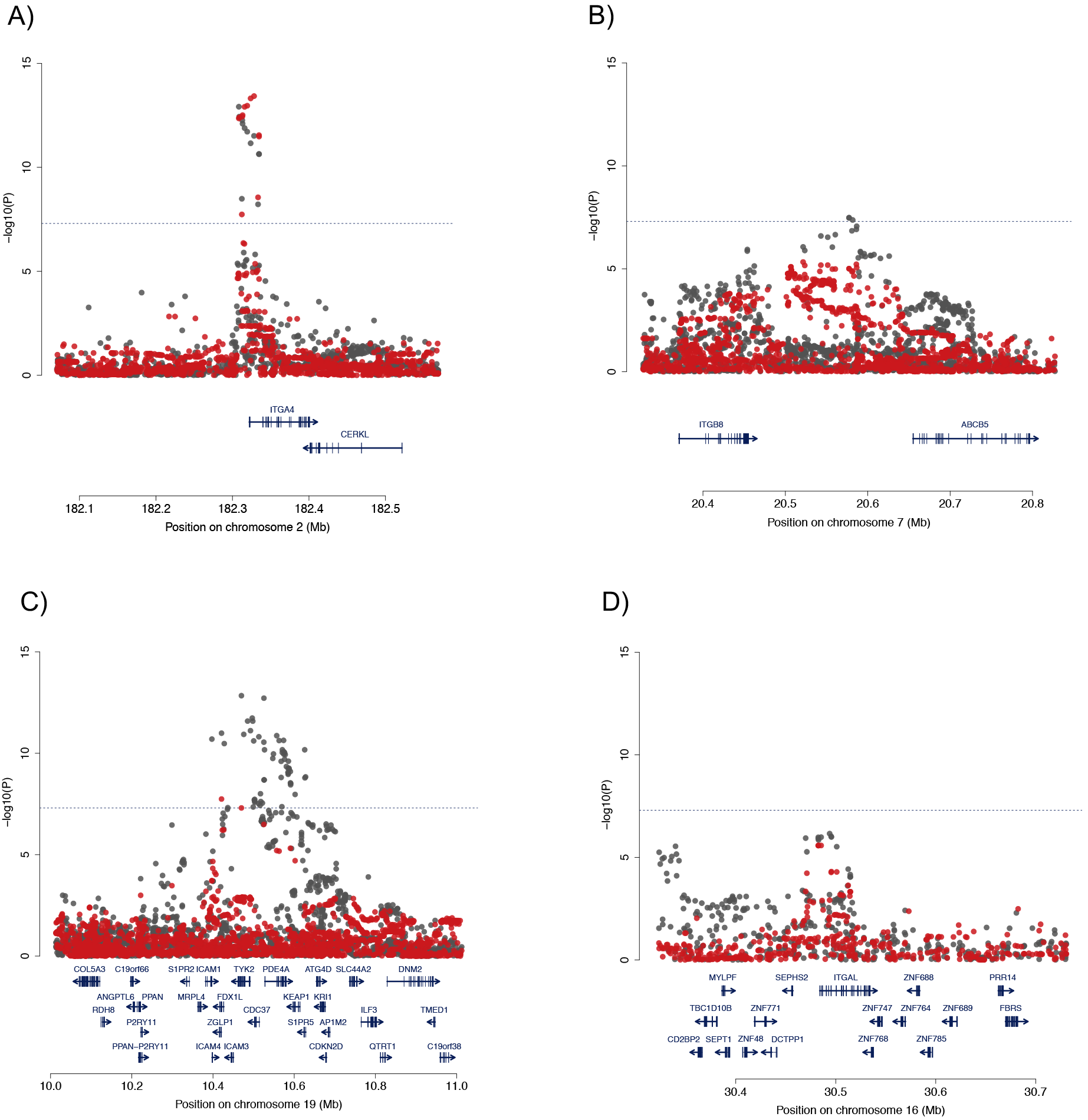
Co-localization of disease association and stimulus response eQTLs in monocytes. The local pattern of disease association (IBD: (A) ***ITGA4***, (B) ***ITGB8***, (C) ***ICAM1***; (D) UC: ***ITGAL***) in grey, and the association of that variant with response to LPS (lipopolysaccharide) stimulation in red. Evidence of co-localization (probability > 70%) is observed for all for signals.

Integrins and their counter-receptors have recently emerged as important therapeutic targets in IBD. Most promisingly monoclonal antibodies that target the components of the a4p7 dimer, responsible for the gut-homing specificity of leukocyte subsets, have demonstrated efficacy in IBD^28–30^. Additionally, an antisense oligonucleotide targeting ***ICAM1*** has shown promise in the treatment of ulcerative colitis and pouchitis^31^. The importance of gut-selectivity for therapeutic approaches is highlighted by the success of antibodies that bind the αL and α4 integrin subunits. Whilst αL-directed therapy with efalizumab demonstrated potential in Crohn’s disease^32^, and α4-directed therapy (which binds a4p1 in addition to a4p7 integrin) with natalizumab is licensed in the USA for Crohn’s disease^33^, both medications have been associated with occurrences of progressive multifocal leukoencephalopathy (PML). This potentially fatal condition is likely mediated by impaired leukocyte migration to the central nervous system leading to JC virus infection of the brain. Owing to the risk of PML, efalizumab has been withdrawn from the market and natalizumab is not licensed for Crohn’s disease in Europe.

Integrins are not only important in cell trafficking, but can also participate in cellular signalling. For example, the αVβ8 heterodimer - both subunits of which are now within confirmed IBD loci - is a potent activator of TGFβ^34^, with a range of cell-type specific effects. Furthermore, mice with dendritic-cell specific deletion of this complex had impaired regulatory T cell function and severe colitis^35^, whereas deleting the complex in regulatory T cells themselves prevented them from suppressing pathogenic T cell responses during active inflammation^36^. While no current IBD therapeutics target αVβ8 directly, promising early results of an oral antisense oligonucleotide to the inhibitory TGFβ-signalling protein SMAD7^37^, itself encoded by a locus identified by genetic association studies^23^, demonstrate the therapeutic potential of modifying TGFβ signaling in Crohn’s disease.

In addition to the connections to anti-integrin and anti-TGFβ therapies described above, IBD GWAS have previously implicated loci containing other therapeutically relevant genes, such as those in pathways targeted by anti-TNF and anti-p40 IBD therapies (**Figure 3**). These discoveries have demonstrated that the importance of the biological pathways underlying associations, and their potential therapeutic relevance, are not necessarily reflected in their GWAS effect sizes. For example, the modest odds ratios of the signals near integrin genes (1.10-1.12) required tens of thousands of samples to detect at genome-wide significance. Furthermore, analyses aimed at understanding the specific cellular contexts in which these genes are active in IBD, as well as the risk-increasing direction of effect (e.g. consistent up-regulation of integrins in response to LPS stimulus), are only beginning to bear fruit.

**Figure 3.**
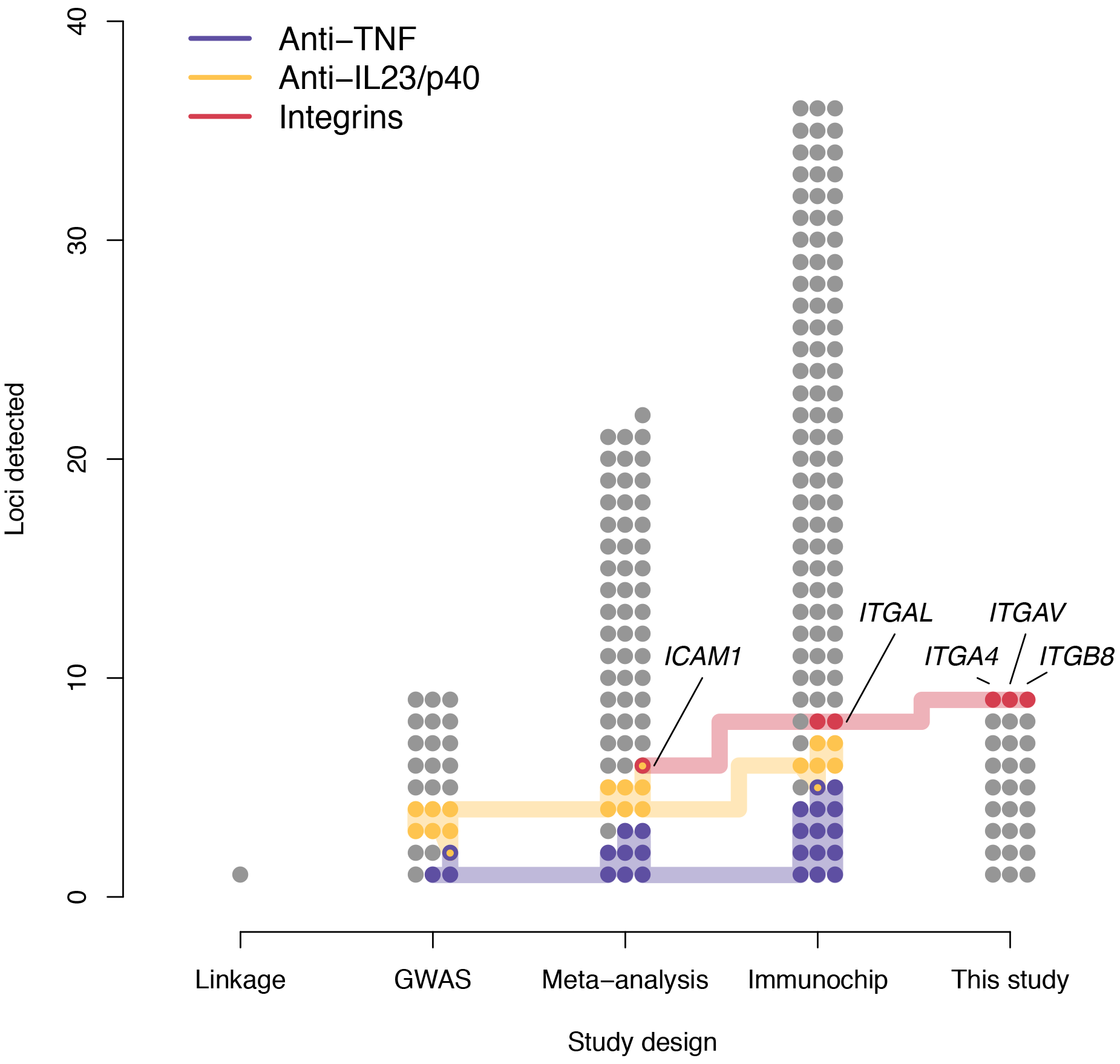
IBD-associated loci containing genes related to known drug targets. All IBD loci are divided into the studies where they were first identified^1^. The direct targets and closely related genes of three classes of IBD therapeutics are highlighted, with particular focus on the newly discovered integrin associations. Despite the general pattern that effect size decreases from left to right, therapeutically relevant associations continue to be found.

Our study has demonstrated that continuing to pursue GWAS, even in a well studied complex disease like IBD, has the potential to complement other powerful approaches, such as targeted genotyping (via the Immunochip) and large-scale genome and exome sequencing. In two cases we have implicated genes in which different variants have previously been shown to cause immune-related Mendelian disorders, echoing a connection made to the very first Crohn’s disease risk gene, ***NOD2***, in which rare missense mutations cause the autosomal dominant granulomatous disorder Blau syndrome^38^. Finally, while the individual effect sizes of our newly discovered associations are modest, we believe that our results show that GWAS continues to deliver new understanding of disease biology and new therapeutic opportunities.

## Acknowledgements

We would like to thank all individuals who contributed samples to the study. This work was cofunded by the Wellcome Trust [098051] and the Medical Research Council, UK [MR/J00314X/1]. Case collections were supported by Crohn’s and Colitis UK. KMdL, LM, CAL, YL, DR, JG-A, NJP, CAA and JCB are supported by the Wellcome Trust [098051; 093885/Z/10/Z; 094491/Z/10/Z]. KMdL is supported by a Woolf Fisher Trust scholarship. CAL is a clinical lecturer funded by the NIHR. We thank Anna Stanton for co-ordinating the Guy’s and St Thomas’ patient recruitment. We acknowledge support from the Department of Health via the NIHR comprehensive Biomedical Research Centre awards to Guy’s and St Thomas’ NHS Foundation Trust in partnership with King’s College London and to Addenbrooke’s Hospital, Cambridge in partnership with the University of Cambridge. This research was also supported by the NIHR Newcastle Biomedical Research Centre.

## Author contributions

KMdL, LM, YL, LJ, DLR, CAA, and SGJ performed statistical analysis. KMdL, LM, YL, LJ, JCL, JGA, SGJ, CAL, NAK, and CAA analysed the data. GH, ERN, CE, CM, AS, DCW, MT, AH, CGM, MP, WGN, CWL, HU, CH, NJP, TA, JCM, JackS, JerS, and PH contributed samples/materials. CAA, JCB, KMdL, LM, JCL, CGM, MP, CAL, NAK, YL, and PH wrote the paper. JCB, CAA, JCM, MP, CWL, TA, and NJP conceived & designed experiments. JCB and CAA jointly supervised research. KMdL, LM, and JCL contributed equally to this work.

## Online Methods

### New genome-wide genetic data

*GW4S samples and genotyping*. 11,768 British IBD cases, diagnosed using accepted endoscopic, histopathological and radiological criteria, were genotyped on the Human Core Exome v12.1. 10,484 population control samples genotyped on the Human Core Exome v12.0 were obtained from the Understanding Society Project. Genotypes were called using optiCall^39^.

*GW4S quality control*. We removed variants that did not overlap between the two versions of the chip, had missingness > 5%, a significant difference in call rate between cases and controls (P < 1×10^−5^), deviated from Hardy-Weinberg equilibrium (HWE) in controls (P < 1×10^−5^), or that were affected by a genotyping batch effect (significant association [P < 1×10^−5^] between an outlier group of cases discovered using principal component analysis [PC1 < −0.005], and the remainder of the samples). We then removed samples with missingness > 1%, heterozygosity ±3 standard deviations from the mean, mismatch between reported and genotypic sex, first-degree relatives or closer (kinship coefficient > 0.177), and non-European samples identified through principal component analysis with HapMap3 populations. After quality control, data were available for 4,474 Crohn’s disease, 4,173 ulcerative colitis, 592 IBD-unclassified cases and 9,500 controls for 296,203 variants.

*Whole-genome sequenced samples*. We generated low-coverage whole genome sequences for 4,686 IBD cases and 3,781 population controls from the UK IBD Genetics Consortium (UKIBDGC) and UK10K Consortium, respectively. Detailed information on sequencing, genotype refinement and quality control are described elsewhere^8^.

*Imputation*. These sequences were combined with 2,504 samples from the Phase 3 v5 release of the 1000 Genomes project (2013-05-02 sequence freeze) to create a phased imputation reference panel enriched in IBD-associated variants. We used PBWT^40^ to impute from this reference panel (114.2 million total variants) into our new GWAS described above.

### Association testing, meta-analysis, and quality control

*Association testing*. Prior to association testing, we removed all samples that were included in previous IBD GWAS meta-analyses (Supplementary Table 1). We then tested for association to ulcerative colitis, Crohn’s disease and IBD separately within the sequenced samples and new GWAS using SNPTEST v2.5, performing an additive frequentist association test conditioned on the first ten principal components for each cohort. We filtered out variants with minor allele frequency (MAF) < 0.1%, INFO < 0.4, or strong evidence for deviations from HWE in controls (p_HWE_<1x10^−7^).

*Meta-analysis*. We used METAL (release 2011-03-05) to perform a standard error weighted metaanalysis of our sequencing and GWAS cohorts with the publicly available International Inflammatory Bowel Disease Genetics Consortium (IIBDGC) meta-analysis summary statistics^1^, after applying the additional MAF ≥0.1%, and INFO ≥0.4 filters to the IIBDGC data.

*Quality control*. The output of the fixed-effects meta-analysis was further filtered, and sites with high evidence for heterogeneity (I^2^>0.90) were discarded. Only sites for which all cohorts passed our quality control filters were included in our analysis. In addition, we discarded genome-wide significant variants for which the meta-analysis p-value was not lower than all of the cohort-specific p-values.

*LD score regression*. We performed LD score regression using LDSC v1.0.0 and European linkage disequilibrium (LD) scores from the 1000 Genomes Project (downloaded from https://data.broadinstitute.org/alkesgroup/LDSCORE/eur_w_ld_chr.tar.bz2) on our filtered meta-analysis summary statistics for all sites with INFO > 0.95. This INFO threshold is to avoid confounding due to poor imputation, as recommended by the authors^41^.

### Locus definition

*Computing LD windows*. An LD window was calculated for every genome-wide significant variant in any of the three traits (Crohn’s disease, ulcerative colitis, IBD), defined by the left-most and right-most variants that are correlated with the main variant with an r^2^ of 0.6 or more. The LD was calculated in the GBR and CEU samples from the 1000 Genomes Phase 3, release v5 (based on 20130502 sequence freeze and alignments). Loci with overlapping LD windows, as well as loci whose lead variants were separated by 500kb or less, were subsequently merged, and the variant with the strongest evidence of being associated was kept as the lead variant for each merged locus.

*Identifying novel loci*. A locus was annotated as known if it contained at least one variant previously reported at genome-wide significance (irrespective of the LD between that variant and the most associated variants in the locus). To ensure that putatively novel signals were not due to long-range LD with variants in previously reported loci, we conducted conditional analysis in our new GWAS for all variants in loci which were less than 3Mb away from a known locus. Putatively novel loci already known in a lower order IBD trait (e.g. a previously known Crohn’s disease locus coming up as an IBD locus) were also removed from this list. This did not apply where, for example, a known Crohn’s disease locus was now associated with ulcerative colitis, or vice versa.

### Fine-mapping

Approximate Bayes factors were calculated from the meta-analysis effect sizes and standard errors described above by applying equation (2) of Wakefield^42^, assuming a prior variance on the log odds ratios of 0.04 (the default prior used by the software SNPTest, and used by Maller *et al*^43^). We then performed fine-mapping using these Bayes factors as described in Maller et al to calculate the posterior that each variant is causal, and the 95% credible set for each association (the smallest set of variants with posteriors that sum to at least 95%). For each association we use the meta-analysis results for the phenotype (Crohn’s disease, ulcerative colitis or IBD) specified in Table 1. We only consider a locus to be confidently fine-mapped if there are no variants in the Phase 3 v5 release of the 1000 Genomes project (2013-05-02 sequence freeze) in high LD (r^2^ ≥ 0.6) with our hit SNP, but missing from our dataset, and no variants in our data within high LD (r^2^ > 0.8) that fail during our QC procedure.

### eQTL overlap

*Identifying eQTL overlaps*. Nine eQTL datasets were searched to identify variants within the 26 newly identified IBD risk loci that are associated with variation in gene expression (Supplementary Table 5). Splice-QTLs based on exon-ratio^44^ and transcript-ratio^45–47^ were also included in the search where available (Supplementary Table 5). The most significant variant-gene associations were extracted from each eQTL/splice-QTL dataset and were reported as candidates if that variant had r2 > 0.8 with any of the lead SNPs in the 26 IBD risk loci.

*Testing for co-localization*. We tested for co-localization between IBD association signals and eQTLs using the coloc2 method^48^, implemented in the R package coloc. We used a window size of 250kb on
either side of the IBD association, and implemented the default settings as recommended. Each test was repeated using two different values for the prior probability of co-localization, p_12_: 1×10^−5^ and 1×10^−6^.

